# Direct RNA sequencing of Respiratory Syncytial Virus infected human cells generates a detailed overview of RSV polycistronic mRNA and transcript abundance

**DOI:** 10.1101/2021.12.23.473996

**Authors:** I’ah Donovan-Banfield, Sophie Hall, Tianyi Gao, Eleanor Murphy, Jack Li, Ghada T. Shawli, Julian Hiscox, Rachel Fearns, David A. Matthews

## Abstract

To characterize species of viral mRNA transcripts generated during respiratory syncytial virus (RSV) infection, human fibroblast-like MRC5 lung cells were infected with subgroup A RSV for 6, 16 and 24 hours. Total RNA was harvested and polyadenylated mRNA was enriched and sequenced by direct RNA sequencing on an Oxford nanopore device. This yielded over 150,000 direct mRNA transcript reads which were mapped to the viral genome and analysed to determine relative mRNA levels of viral genes using our in-house ORF-centric pipeline. We were also able to examine frequencies with which polycistronic readthrough mRNAs were generated and to assess the length of the polyadenylated tails for each group of transcripts. We show that there is a general but non-linear decline in gene transcript abundance across the viral genome, as predicted by the model of RSV gene transcription. However, the decline in transcript abundance is not consistent. We show that the polyadenylate tails generated by the viral polymerase are similar in length to those generated by the host cells polyadenylation machinery and broadly declined in length for most transcripts as infection progressed. Finally, we observed that the steady state abundance of transcripts with very short polyadenylate tails is much less for N, SH and G transcripts compared to NS1, NS2, P, M, F and M2 which may reflect differences in mRNA stability and/or translation rates.

## Introduction

Respiratory syncytial virus (RSV) causes a respiratory infection that leads to significant levels of morbidities and mortalities in infants and young children across the globe (1, 2). Recovery from infection does not lead to long term protection and repeat reinfections over an individual’s lifetime are a hallmark of RSV (3, 4). Thus, hospital admission of elderly patients with life-threatening complications of RSV infection are also common (5, 6). Typically, RSV represents a global viral respiratory disease burden comparable to that of influenza virus, which underlines the importance of trying to understand this virus’ lifecycle (7–9).

The RSV genome is single stranded, negative sense RNA approximately 15,000 nt in length. The genome codes for 10 major capped and polyadenylated mRNAs in the order *NS1, NS2, N, P, M, SH, G, F, M2* and *L* (Figure 1A) (10–12). In virus particles and infected cells, the viral genome exists as a nucleocapsid structure in which the RNA is coated along its length by the nucleocapsid protein (N), and associated with a viral RNA dependent RNA polymerase known as L, a phosphoprotein (P) and viral protein M2-1 (13, 14). Transcription of the RSV genome depends on the polymerase complex recognizing and responding to *cis-acting* elements within the viral genome. Each of the genes is flanked with conserved elements referred to as gene start (GS) and gene end (GE) signals (12). Most of the genes are separated by short intergenic regions, although there is one overlapping gene junction, in which the GS signal of the downstream L gene lies 48 nt upstream of the GE signal of the preceding M2 gene (15).

**Figure 1.**
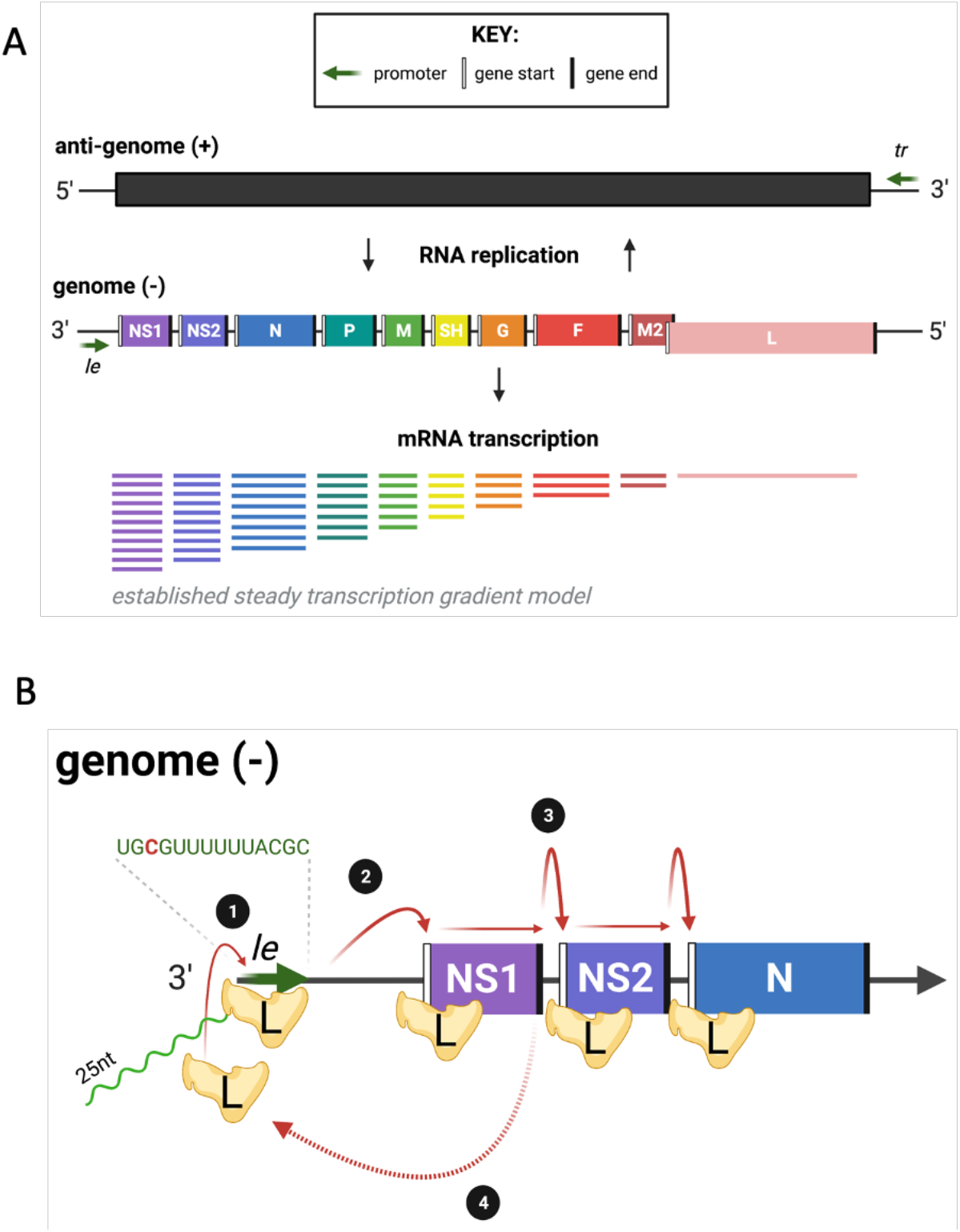
RSV genome and mRNA transcription. (A) A schematic of RSV genome organisation, replication and the mRNA transcripts generated by a simple gradient of transcript initiation. *Le*, leader; *tr*, trailer. (B) A zoomed in simplified schematic of the viral RdRp, L, binding to the negative-sense genome to transcribe viral mRNA. L binds to the *le* promoter (1) and initiates transcription opposite position 3 of the Le promoter, generating a ~20-25 nt transcript (shown in green) which is released before the polymerase scans for the *NS1* gene start signal (vertical white line) (2) from here the mRNA is transcribed until it reaches a gene end signal (vertical black line) and poly-adenylates the nascent mRNA. L will then either continue scanning along to the next gene start signal (3) or fall off and have to re-associate with the genome at the le promoter and start RNA synthesis again (4).

The 3’ end of the genome contains a promoter referred to as the leader or Le promoter. Studies examining the effect of UV exposure on RSV gene expression showed that genes proximal to the 3’ end of the genome were more resistant to UV damage than genes from the 5’ end of the genome (16). This finding established that RSV genes are transcribed sequentially from the 3’ to the 5’ end of the genome. Subsequent minigenome studies confirmed that transcription of a downstream gene is dependent on transcription of an upstream gene, and that none of the intergenic regions contains a promoter (17–19). Based on these and other studies, the prevailing model for RSV transcription is that the polymerase initiates each cycle of transcription at the Le promoter. Subsequent studies showed that the polymerase initiates transcription opposite position 3C of the Le promoter and generates and then releases a short, heterogeneous RNA transcript of ~ 20-25 nt in length. It then scans to locate the GS signal of the first gene where it reinitiates RNA synthesis and caps the mRNA. The polymerase then elongates the mRNA until it reaches a GE signal where it polyadenylates the mRNA by a reiterative stuttering mechanism, and then releases it (Figure 1B) (20, 21). Occasionally, the polymerase fails to recognize a GE signal, resulting in synthesis of a polycistronic mRNA. The possibility also exists that the polymerase could fail to re-initiate RNA synthesis at a GS signal. It is thought that in this case, the polymerase dissociates from the viral genome ending that cycle of mRNA transcription. Because initiation of a new cycle of transcription depends on the polymerase associating with the Le promoter, a transcription gradient is established with genes that are proximal to the 3’ end of the genome molecule being transcribed more than those nearer the 5’ end.

Direct evidence for a transcription gradient was originally obtained by quantifying mRNAs generated in an *in vitro* transcription system and from studies using a small molecule polymerase inhibitor that prevented initiation at the promoter (22, 23). However, there have been several attempts to quantify the relative expression of RSV mRNA which do not fully agree with these experiments. Some of these studies have involved Illumina based RNA-seq analysis of intracellular viral RNA transcripts, which showed a deviation from the expected transcription gradient (24). In another study, RT-qPCR analysis of mRNA levels in RSV infected cells revealed genotype-specific variations in mRNA levels, but with *G* having a higher mRNA level than *N* in each genotype tested, a finding that could not be accounted for by RNA stability (25). Based on these findings, it was proposed that in some cases the polymerase might scan through upstream genes in a non-transcribing mode and then initiate at a downstream GS signal, and/or that some gene junction signals facilitate polymerase recycling so that certain genes are transcribed reiteratively (25). In this model, after reaching the GE signal for the G gene, for example, the polymerase might scan upstream to re-initiate mRNA transcription at the G gene GS signal, this would entail almost 1000 nt of upstream scanning. In support of this it has been shown that the polymerase can scan upstream from the M2 GE signal to locate the GS signal for the L gene (18) but this is a relatively small distance of less than 68 nt (in the case of RSV-A). However, these recent studies suffer to one degree or another from confounding factors. For example, reverse transcription and PCR amplification, which are necessary steps in Illumina based RNA-seq experiments, could be variable in efficiency depending on the mRNA transcript sequence and secondary structure. Likewise, RT-qPCR based quantification of different viral genes could be influenced by the selection of standards, used to determine copy number, that do not accurately mimic viral mRNAs.

Direct RNA sequencing (dRNAseq) has been used to study transcriptomes from a range of sources including human and viral transcriptomes (26–31). The technique sequences mRNA directly from the polyadenylated tail towards the 5’ cap by feeding the mRNA molecule through a nanopore and measuring changes in current across the pore as nucleotides pass through. Inherently dRNAseq can suffer from a 3’ bias as it is not possible to determine if the molecule sequenced is indeed full length with an authentic cap or if it is degraded. In addition, there is a significant rate at which errors are introduced into the sequencing data – approximately 10% of nucleotides are wrong or missing. Nonetheless, the nature of the technique offers the opportunity to quantify mRNA abundance directly without potentially confounding experimental steps (e.g. reverse transcription and PCR) and it allows for sequencing of the whole length of the mRNA providing important data on the structure of the mRNA including if the mRNA is polycistronic. Finally, it is also possible to determine poly A tail lengths (32).

We have previously used dRNAseq to examine the transcriptome of human adenoviruses, SARS-CoV-2 and adenovirus vector-based vaccines illustrating the significant advantages this approach offers in a range of different settings (26, 30, 33). Here we apply the technique to examine the transcriptomic repertoire of human RSV infection of MRC-5 cells over three time points. We show that there is a gradient of transcription as initially proposed in early studies but that it is not strictly linear. We also provide insights into the rate of production of viral polycistronic mRNA and show that the poly A tails of RSV mRNA are similar in length to those of human mRNA. These findings cement the utility of dRNAseq for analysing viral transcript frequencies.

## Materials and Methods

### Virus and cells

Human MRC-5 cells, a genetically normal human lung fibroblast-like line, were obtained from European Collection of Authenticated Cell Cultures (#05072101, ECACC). The cells were cultured in DMEM supplemented with 10% foetal bovine serum, 100 U/ml penicillin and 100 μg/ml streptomycin. After reaching confluence, the cells were infected with RSV strain A2 at a multiplicity of 10. The infected cells were harvested at 6, 16 and 24 hours post infection.

### RNA extraction and sequencing

RNA extraction and sequencing was as described previously(26, 30, 33). Briefly, total RNA was extracted from the infected cell using TRIzol reagent (#15596026, Ambion) at 1ml of reagent per 10^7^ cells and the RNA extracted as per manufacturer’s recommendations except that the final wash of the RNA pellet in 70% ethanol was repeated a further two times (total of 3 washes). Total RNA was enriched for polyA tails using Dynabeads™ mRNA purification kit (#61006, Invitrogen) as per manufacturer’s instructions. We used the SQK-RNA002 kits and MIN106D R9 version flow cells (Oxford Nanopore Technologies), following the manufacturer’s protocols.

### Data analysis, mapping to viral genome and human transcripts

Reads were mapped to a fasta file containing the RSV A2 genome (accession number KT992094.1) using minipmap2(34) (command line: minimap2 -a -x map-ont -uf -k14 --sam-hit-only RSV_AB.fasta RSV2020_16hpi.fastq > map_2_RSV.sam). Alternatively we mapped the reads to a list of human cDNA transcripts as previously described (command line: minimap2 -a -x map-ont -uf -k14 --sam-hit-only H_sapiens.GRCh38.cdna.fasta RSVA2_24hpi.fastq > mapped_2_HSap_cDNA.sam) (33).

### Data analysis, characterisation of viral transcripts

To analyse nanopore transcripts that have a very high error rate we developed a software pipeline that assigns transcripts according to the ORFS the code(26). Briefly, the pipeline takes the mapping information form the SAM file to group the transcripts mapping to the viral genome according to shared start locations, exons (or equivalent) and stop locations. Once grouped in this manner, a representative transcript is then generated for each transcript group based solely on the genome sequence. User supplied information on the location of known ORFs is employed to assign transcript groups according to what known ORFS would be coded by each transcript group. In this way the pipeline determines the percentage of mRNAs that express each ORF relative to the total number of transcripts that map to the target genome. This pipeline generates tables describing the structure of transcripts, what ORFs are present and how many mRNA molecules belong to each group of transcripts. Allied to this analysis, nanopolish (35) was used to determine the polyA length of each sequenced transcript and in house scripts grouped transcripts according to GE usage before compiling a list of polyA lengths for generating violin plots. For the ORF centric pipeline analysis, only transcripts that were QC-flagged as PASS by nanopolish and had an estimated polyA tail length of 20 or more were considered (approximately 60% of transcripts). For the analysis involving short polyA lengths, again only transcripts QC-flagged as PASS by nanopolish were considered. Visualisation of differences in polyA tail lengths of transcripts sharing the same GE signal was conducted in R 4.1.2 using RStudio, using ggplot2 (v3.3.5) and tidyverse (v1.3.1) packages.

### Data analysis of G glycoprotein transcripts

Previously, and in line with other groups using dRNAseq, we had noted that the 5’ most 10 or so nucleotides of an mRNA molecule are lost during nanopore sequencing (26, 27). To compensate for this in our analysis pipeline we use the genome sequence to programmatically add back the missing 10 nucleotides upstream and this approach works well for transcripts where the authentic initiating methionine is close to the 5’ cap. In the case of the G protein of RSV however this approach leads the software to utilise an out of frame AUG some 8 nt upstream of the authentic AUG. The software pipeline thus reports a large number of transcripts in which the apparent primary ORF is short and unknown but is immediately followed by a second ORF which is correctly identified as the G protein. For the purposes of this analysis we have counted these as genuine G gene transcripts since manual inspection of the structure of these transcripts is consistent with the GS for G protein mRNA.

### Data Availability

Raw fastq files, associated fasta files and polyA length output files are available from Zenodo, DOI: **https://doi.org/10.5281/zenodo.5799655**

### Code Availability

The software is available from the authors on request or from Zenodo, DOI: **http://doi.org/10.5281/zenodo.3610249**

## Results

### Overall map of RSV transcripts

To examine RSV transcripts using dRNAseq, MRC-5 cells were infected with RSV strain A2 at an moi of 10 pfu/ ml. Cells were then harvested at 6, 16 and 24 hpi to encompass the approximate length of time for a single cycle of virus replication. RNA was isolated and subjected to dRNAseq. Table 1 lists the number of sequence reads from each timepoint and the number of reads that map to the RSV genome or the human transcriptome in each sample. The number of reads mapped to the RSV genome climbed dramatically over the course of infection, from just over 3,000 reads at 6 hours post infection to over 147,000 reads by 24 hours post infection.

**Table 1.** Read counts and mapping information. Table to show total number of reads, longest read and average read length for each dataset alongside mapping data for reads mapping to human transcripts or the RSV genome. For the FASTQ data we also indicate the longest read length and average read length. For the mapped data we also indicate the length of the longest read mapped.

| Sample | FASTQ count (Longest read : Average length) | Mapped to RSV (Longest mapped read) | Mapped to Human Transcripts (Longest mapped read) |
| --- | --- | --- | --- |
| **MRC5 mock** | 2,195,978 (16,860 : 1358.5) | 0 | 1,945,497 (16,859) |
| **MRC5 6hpi** | 1,577,904 (22,167 : 1426.4) | 3192 (8,015) | 1,328,890 (22,166) |
| **MRC5 16hpi** | 2,380,322 (19,372 : 1367.1) | 76,702 (9,880) | 1,992,491 (19,371) |
| **MRC5 24hpi** | 1,585,580 (12,629 : 1145.4) | 147,323 (7,196) | 1,125,920 (12,628) |

Broadly, the pattern of mapped reads shows a gradient of transcription, with genes that are proximal to the 3’ Le promoter being more highly represented (Figure 2). In addition, we can also see that, as a result of the sequencing technology which sequences transcripts polyA tail first, there is greater read depth within each gene nearer the location of the polyA site. The relative levels of mRNA were somewhat different at 6 hpi than at 16 and 24 hpi, with *NS1* and *NS2* transcripts being more highly represented at 6 hpi than at later times. This could be due to asynchronous viral entry, such that in some cells, only the first two genes had been transcribed by 6 hpi. We also cannot exclude the possibility that this early timepoint also represents mRNAs that were packaged into virus particles, rather than those that were synthesized in the infected cells. The read patterns at 16 and 24 hpi are similar to each other.

**Figure 2.**
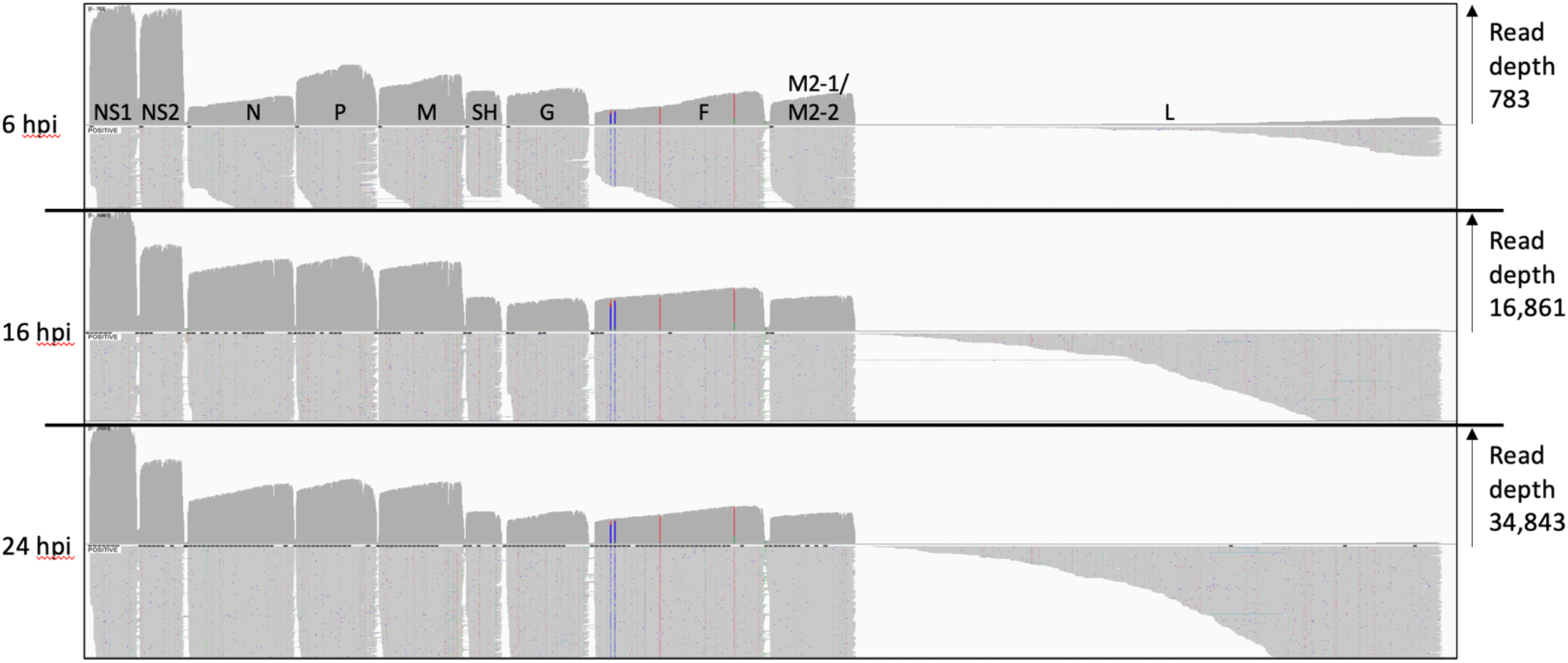
Reads mapped to the RSV genome. An IGV generated image of the dRNAseq reads mapped to the viral genome at 6, 16 and 24 hours post infection, the location of the viral ORFs is indicated above the top panel. Each panel shows the overall depth of reads along the genome in the top half and the bottom half shows a selection of the mapped reads illustrating how they align to the viral genome. The vertical coloured lines within the F ORF pinpoint three SNPs present in the observed reads relative to the reference genome for RSV A2.

We used our previously described ORF centric analysis pipeline (26) to examine the number of mRNA molecules that code for a known gene, and Table 2 shows the relative contribution of mRNA that code for each RSV gene. This data is a subset of the total mapped reads shown in figure 2 in that it only includes dRNAseq reads with a polyA tail of over 20 nt. This 20 nt cut-off is used so that we can be highly confident that the transcripts being counted are genuine mRNA molecules. Notably this table indicates that whilst *NS1* and *NS2* dominate at 6 hours post infection, only *NS1* seems to dominate thereafter as levels of *NS2* gene expression are then closer to *N* gene expression. Moreover, rather than a gene-by-gene step-wise decline in mRNA abundance the more distal the gene is from the 3’ Le promoter there is instead perhaps four groups of gene abundance by 16 and 24 hours post infection (Table 2 – see abundance relative to NS1 transcripts data). Group 1 containing just NS1, group 2 comprising *NS2*, *N*, *P* and *M* (around half the abundance of NS1) followed by group 3 with *SH, G, F* and *M2-1/M2-2* (each around 20% of the abundance of NS1) with group 4 containing just L. This matches the visual pattern of depth of aligned transcripts shown in figure 2 and these data are consistent with a model of obligatorily sequential transcription. As with our previous publications using dRNAseq (26), there is a large number of transcripts which could not be confidently associated with a particular gene (listed as “not identified”) these usually represent truncated RNA molecules which arise from mRNA degradation, mechanical shearing or molecules with apparent short deletions within the body of the transcript.

**Table 2.** List of ORFs from RSV and their percent contribution to the RSV derived transcriptome. This table illustrates the number of reads mapped that contain the indicated RSV ORFs as the 5’ most ORF. At each timepoint the raw number of individual transcripts is indicated alongside the percent contribution of those transcripts at that timepoint. In addition we have calculated the percent abundance of each transcript relative to NS1 abundance. Note that this table only counts transcripts that have been shown to have a polyA tail over 20nt long.

| Feature | Gene order on viral genome | 6hpi raw count | 6hpi percent of total (percent of NS1) | 16hpi raw count | 16hpi percent of total (percent of NS1) | 24hpi raw count | 24hpi percent of total (percent of NS1) |
| --- | --- | --- | --- | --- | --- | --- | --- |
| Not identified | N/A | 563 | 26.61 | 8,639 | 17.26 | 19,587 | 19.82 |
| NS1 | 1 | 406 | 19.19 | 9,078 | 18.13 | 18,926 | 19.16 |
| NS2 | 2 | 323 | 15.26 (80) | 5,123 | 10.23 (56) | 10,539 | 10.67 (56) |
| N | 3 | 75 | 3.54 (18) | 5,308 | 10.60 (58) | 8,604 | 8.71 (45) |
| P | 4 | 201 | 9.50 (50) | 5,537 | 11.06 (61) | 10,111 | 10.23 (53) |
| M | 5 | 139 | 6.57 (34) | 4,823 | 9.63 (53) | 8,630 | 8.73 (45) |
| SH | 6 | 85 | 4.02 (21) | 2,010 | 4.01 (22) | 4,126 | 4.18 (22) |
| G (initiation at 2nd AUG) | 7 | 82 | 3.88 (20) | 1,559 | 3.11 (17) | 3,203 | 3.24 (17) |
| F | 8 | 49 | 2.32 (12) | 2,447 | 4.89 (27) | 3,850 | 3.90 (20) |
| M2-1 | 9 | 74 | 3.50 (18) | 2,412 | 4.82 (27) | 4,329 | 4.38 (23) |
| L | 10 | 0 | 0 | 5 | 0.01 (0) | 6 | 0.01 (0) |
| Total of all features |  | 2,116 |  | 50,064 |  | 98,801 |  |

### Polycistronic reads

Previous studies in which RSV mRNAs were analysed by Northern blot analysis had revealed that polycistronic species are synthesized in addition to monocistronic mRNAs (11). Polycistronic transcripts are generated when the polymerase fails to recognize a GE signal and continues transcribing into the adjacent downstream gene. The dRNAseq data allows a quantification of the abundance of different polycistronic mRNAs. To visually represent polycistronic messages we used an in-house script to group the alignments according to the mapped location of the 3’ end of individual reads (Figure 3) – the nature of dRNAseq means there is greater confidence that the 3’ end of a molecule has been correctly captured. From this data we can clearly see that in many cases there are significant numbers of polycistronic messages, with three or even four genes on the same mRNA molecule in some cases, consistent with the previously published Northern blot data. To provide an estimate of the relative rates that a GE signal is ignored to enable the production of a polycistronic message we divided the maximum read depth of the intergenic region by the maximum read depth of the preceding gene (Table 3). The proportion of readthrough at each intergenic region was consistent over time, except for higher levels of readthrough at the *NS1-NS2* and *G*-*F* intergenic regions at 6 hpi compared to 16 hpi. We note that there appear to be three broad groups of GE signals, one is characterised by the *NS1*, *M* and *F* GE signals with a relatively high rate of readthrough (9-13%), the second is characterised by the *NS2*, *N*, *P* and *G* GE signals with a more moderate rate of readthrough (2 – 4%) with the SH gene GE signal being the most effective at signalling mRNA release (only 0.5% readthrough).

**Figure 3.**
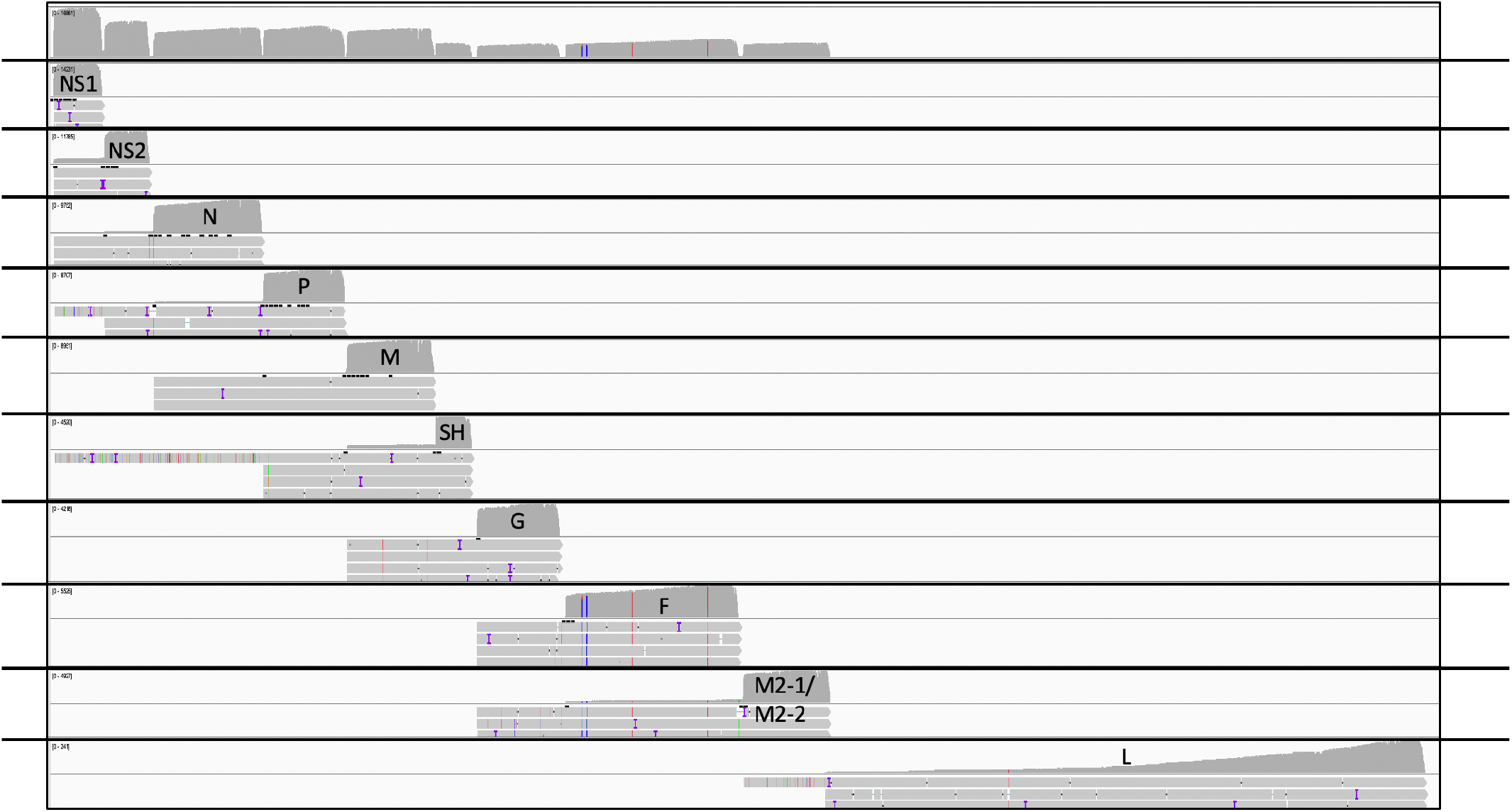
Reads mapped to the RSV genome grouped by GE site usage. An IGV generated image of the dRNAseq reads mapped to the viral genome at 24 hours post infection alongside a series of panels where, in each panel, the reads have been collated according to which GS signal they appear to be utilising. In each panel the top half represents the depth of reads and the bottom half shows a small handful of the longest reads mapping in that group in order to illustrate the presence of polycistronic mRNA in each group.

**Table 3.** Readthrough rates at GE signals. This table shows the percentage of apparent readthrough events within each intergenic region as indicated and listed as the region between each GE and GS signal. This is calculated from the maximum depth of read of the 5’ most gene and the read depth of the intergenic region.

| **GE-GS boundary** | **Percent readthrough at 6hpi** | **Percent readthrough at 16hpi** | **Percent readthrough at 24hpi** |
| --- | --- | --- | --- |
| **NS1-NS2** | 21.6 | 12.9 | 11.9 |
| **NS2-N** | 2.3 | 3.2 | 2 |
| **N-P** | 3.2 | 3.5 | 4 |
| **P-M** | 1.5 | 2.5 | 2.3 |
| **M-SH** | 11.2 | 8 | 8 |
| **SH-G** | 0.5 | 0.6 | 0.5 |
| **G-F** | 0.4 | 2.6 | 2.1 |
| **F-M2** | 10.5 | 9 | 8.6 |

Table 4 shows an analysis of polycistronic mRNA detected at 24 hours post infection. We can see that there is a significant number of polycistronic mRNA produced. This table is calculated from the list of characterised transcripts in supplementary table 3 by counting the number of transcripts that contain one or more genes relative to the number of transcripts that code for just the first gene under consideration. For example, there are 10,324 transcripts containing the *NS2* gene only, compared to 212 *NS2-N* transcripts.

**Table 4.** Abundance of polycistronic messages at 24 hours post infection. This table shows the number of transcripts observed at 24 hours with either one gene (monocistronic) or with increasing numbers of whole additional genes present on individual mRNAs (polycistronic) after the indicated 5’ most gene. For each combination, the number of transcripts cited refers to only transcripts that have that indicated structure.

|  | NS1 | NS2 | N | P | M | SH | G | F | M2 |
| --- | --- | --- | --- | --- | --- | --- | --- | --- | --- |
| **First gene only** | 16,863 | 10,324 | 8,310 | 9,923 | 7,960 | 4,112 | 3,159 | 3,577 | 4,165 |
| **First, Second gene only (%)** | 1,997<br>(11.8%) | 212<br>(2.1%) | 278<br>(3.3%) | 169<br>(1.7%) | 537<br>(6.7%) | 14<br>(0.3%) | 42<br>(1.3%) | 271<br>(7.6%) | 0 |
| **First, Second, third gene only (%)** | 30<br>(0.2%) | 1<br>(0.01%) | 5<br>(0.06%) | 10<br>(0.1%) | 1<br>(0.01%) | 0 | 1<br>(0.03%) | 0 | 0 |
| **First, second, third, fourth gene only (%)** | 3<br>(0.02%) | 0 | 1<br>(0.01%) | 0 | 0 | 0 | 0 | 0 | 0 |

### L transcripts which terminate at the M2 GE

The *M2-L* gene junction is unusual, with the *L* GS signal lying upstream of the *M2* transcript GE signal. Thus, to generate full-length *L* mRNA, the polymerase must ignore the *M2* GE signal. We examined how frequently the polymerase that initiates at the *L* gene start terminates at the *M2* GE versus the *L* GE. We were able to detect just two transcripts that start at nucleotide 8492 (which in this dataset is the start location for authentic L transcripts) but terminate at 8559 (the location of transcription termination for the *M2* transcripts in this dataset). At both 16 and 24 hours post infection there was one transcript in each dataset with a poly A tail greater than 20 nt in length compared to 5 and 6 full length *L* transcripts at 16 and 24 hours respectively. This is likely a very conservative estimate given that we only consider transcripts that were read in full length and so the true number of L transcripts will be underrepresented. For example, the read depth at 24 hours near the 3’ end of the L gene suggests that there may be as many as 240 L transcripts at this time point instead of the 6 we report based on full length reads.

### PolyA tail length

The dRNAseq approach captures data on the length of the polyA tail if one is detected. Figure 4 shows violin plots of the polyA tail length for all the transcripts used to quantitate gene expression (i.e. polyA tail > 20 nt) whose 3’ ends map to the classical dominant GE signals for each gene. For viral mRNA the interquartile range of the polyA tail length is between 50 and 200 nt. In addition, we note that for all transcripts grouped by GE signal, between 16 and 24 hours the median poly A tail length declines (Figure 4; Supplementary tables 1, 2 and 3).

**Figure 4.**
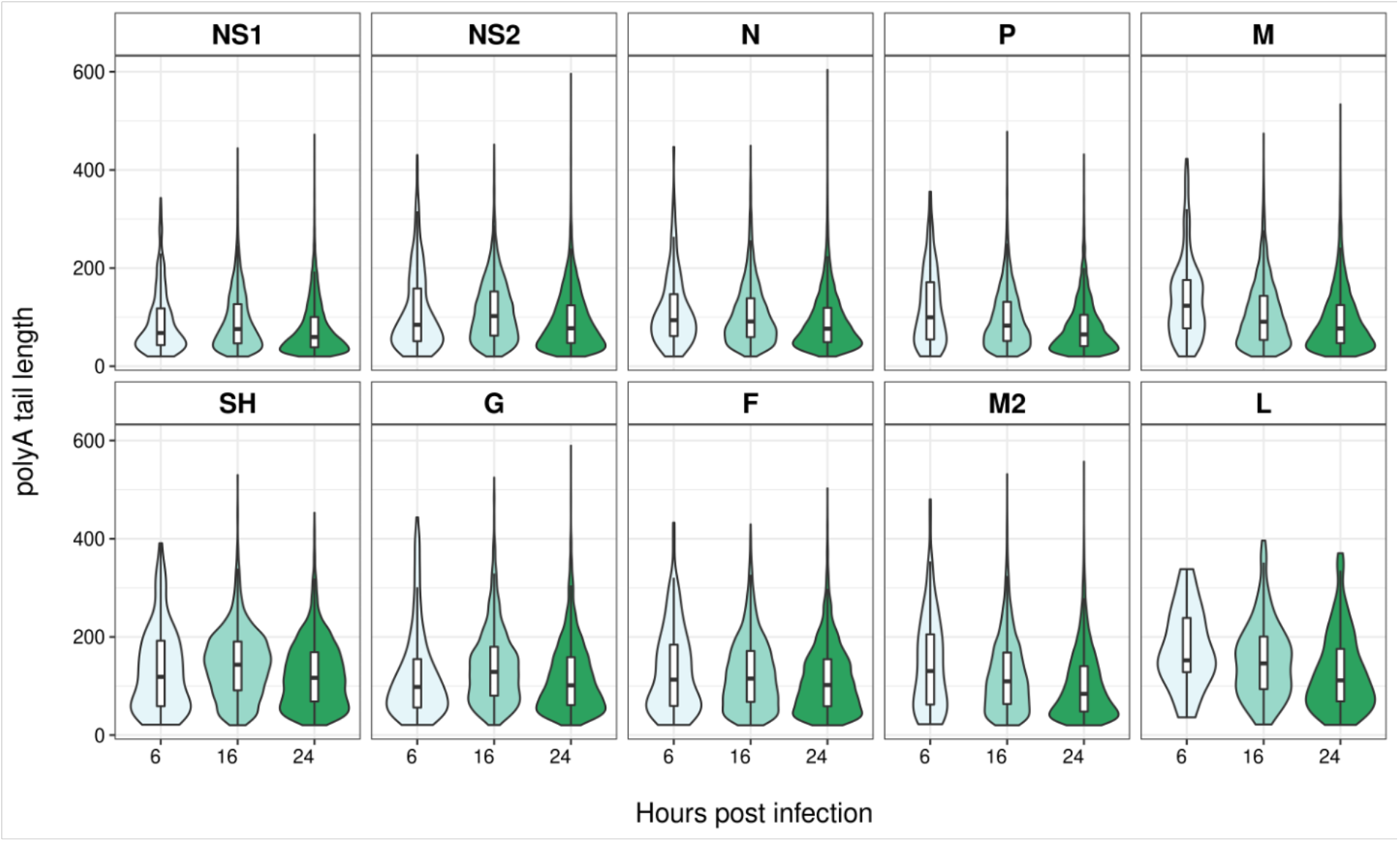
Violin plots of polyA tail lengths of transcripts terminating at different GE sites. A series of violin plots showing the measured polyA tail lengths of transcripts that share the GE signal following each individual ORF on the RSV genome. The different time points are indicated in shades of green from light to dark for increasing hours post infection (6, 16 and 24). In addition, a boxplot is superimposed on each violin plot, showing the median and interquartile range.

Transcripts with very short polyA tails are associated with both inefficient translation and low stability (36). We used an in-house script to identify RSV transcripts from each time point which had short polyA tail lengths (<=20 nt) that nanopolish determined was suitable to estimate a poly A tail length for. When these transcripts were mapped to the RSV genome, we noted that for the most part the distribution of read depth was similar except for *N*, *SH* and *G* transcripts which were notably underrepresented at 16 and 24 hours post infection (Figure 5). To quantify this further we processed the mapped reads with a polyA tail less than 20 nt using our ORF centric pipeline. The data at 6 hours post infection is harder to interpret with confidence due to the low overall read depth. Table 5 provides a more quantitative analysis of the data at 24 hours post infection illustrating the dramatic fall in the relative contributions of *N*, *SH* and *G* transcripts with polyA tails less than or equal to 20 nt in length.

**Figure 5.**
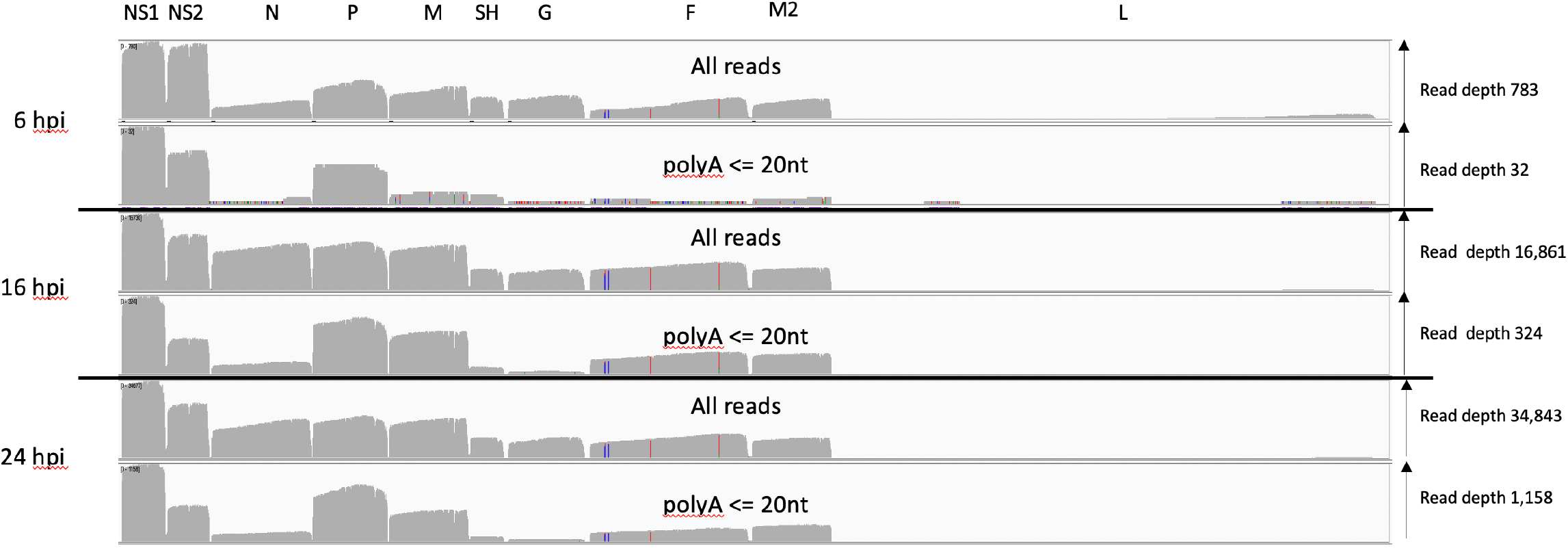
Reads mapped to the RSV genome grouped by polyA tail length. An IGV generated image of the dRNAseq reads mapped to the viral genome split into three sections representing the mRNA sequenced at 6, 16 and 24 hours post infection, the location of the viral ORFs is indicated along the top. Each section has two panels, the top panel shows the overall depth of reads along the genome for all the mapped reads and the lower one for only reads with a polyA length of between 1 and 20 nucleotides as reported by nanopolish software.

**Table 5.** Contribution to the RSV transcriptome for transcripts with short polyA tails at 24 hpi. This table illustrates the number of reads mapped that contain the indicated RSV ORFs as the 5’ most ORF. For each ORF raw counts and percent contribution have been calculated for transcripts with a poly A tail over 20nt and those with a poly A tail equal to or less than 20nt.

| Feature | Gene order on viral genome | poly A <=20 raw count | poly A <=20 percent of total (percent of NS1) | polyA > 20 raw count | polyA > 20 percent of total (percent of NS1) |
| --- | --- | --- | --- | --- | --- |
| **Not identified** | N/A | 607 | 16.94 | 19,587 | 19.82 |
| **NS1** | 1 | 1028 | 28.63 | 18,926 | 19.16 |
| **NS2** | 2 | 369 | 10.30 (36) | 10,539 | 10.67 (56) |
| **N** | 3 | 93 | 2.59 (9) | 8,604 | 8.71 (45) |
| **P** | 4 | 628 | 17.52 (61) | 10,111 | 10.23 (53) |
| **M** | 5 | 344 | 9.60 (33) | 8,630 | 8.73 (45) |
| **SH** | 6 | 36 | 1.00 (4) | 4,126 | 4.18 (22) |
| **G (initiation at 2nd AUG)** | 7 | 26 | 0.73 (3) | 3,203 | 3.24 (17) |
| **F** | 8 | 101 | 2.82 (10) | 3,850 | 3.90 (20) |
| **M2-1** | 9 | 186 | 5.20 (18) | 4,329 | 4.38 (23) |
| **L** | 10 | 0 | 0 | 6 | 0.01 (0) |
| **Total of all features** |  | 3,584 |  | 98,801 |  |

### Variant transcripts

As we have shown in other viral systems there are a large number of variant transcripts reported by our pipeline (supplementary tables 1-3). In the majority of cases, these likely arise from mRNA molecules that are incomplete due to nucleases or mechanical shearing. In some cases, there appear to be micro deletions, which could potentially be an artefact of the sequencing technology or of the mapping algorithms. For example, the most frequently reported minor deletion at 24 hours post infection is at nucleotides 849 to 862 and there is a very A-rich region at this location. Nanopore sequencing is known to have difficulties accurately reporting homopolymeric runs (37). However, there are additional transcripts that appear to result from more substantive skipping events. For example, screening supplementary table 3 for mRNA transcript groups where at least 10 mRNA molecules were observed reveals over 100 different transcripts with small deletions between 9 and 40 nucleotides long. This totals some 4,000 individual polyadenylated mRNA molecules with deletions of at least 9 nucleotides. This includes, for example, a transcript with a 25 nt deletion within the *NS2* gene between nt 849 and 874 on the virus genome, leading to an out of frame truncation of the *NS2* ORF. For this transcript there were 168 distinct mRNA molecules with an average polyA tail length of 95 nucleotides. This same 25 nt deletion was observed in 77 sequenced transcripts at 16 hours post infection (supplementary table 2) with an average poly A tail length of 107 nt. In addition, we also see cases where there are apparently well utilised polyA sites at significant distance from classical GE signals. For example, there are some 459 transcripts that apparently have a polyA tract that begins near or on nt 3090, almost 200 nt away from the dominant polyA site for the *P* gene transcripts at nt 3244. This group of aberrantly polyadenylated transcripts would all still express full length P protein and have an average polyA tail length of 124 nt.

There are also individual transcripts with unusual structures. For example, within the 24 hr post infection dataset there are 59 transcripts that map to the RSV genome with an insertion of over 100 nt (Supplementary table 3). Examination of the top 10 insertions reveals a mixture of RSV sequences, unknown sequences, and apparently one region of a human mRNA. In addition, we mapped the transcripts to concatenated RSV genomes to see if any transcripts mapped across two duplicate genomes. We identified less than 100 transcripts that were each unique but did indeed apparently map across two genomes concatenated together in the same sense (Supplementary datasets 5 and 6).

### Reads mapping to the human transcriptome

In addition to the reads that map to the viral genome we also captured host cell mRNA data. We have previously analysed this kind of data in adenovirus vaccine infected cells to show that it is potentially of some utility (33). However, given the relatively low number of sequence reads in any given experiment there is no clear consensus in the literature about how reliable dRNAseq is for gene expression analysis or which method is most appropriate. Nonetheless, by counting reads mapped to known human transcripts we do observe that some genes known to be highly induced by RSV infection also appear to be highly induced in our datasets (Table 6 and supplementary table 4). This includes for example ISG15, IL6, IFITM1, MX2, OAS2 (38–41).

**Table 6.**
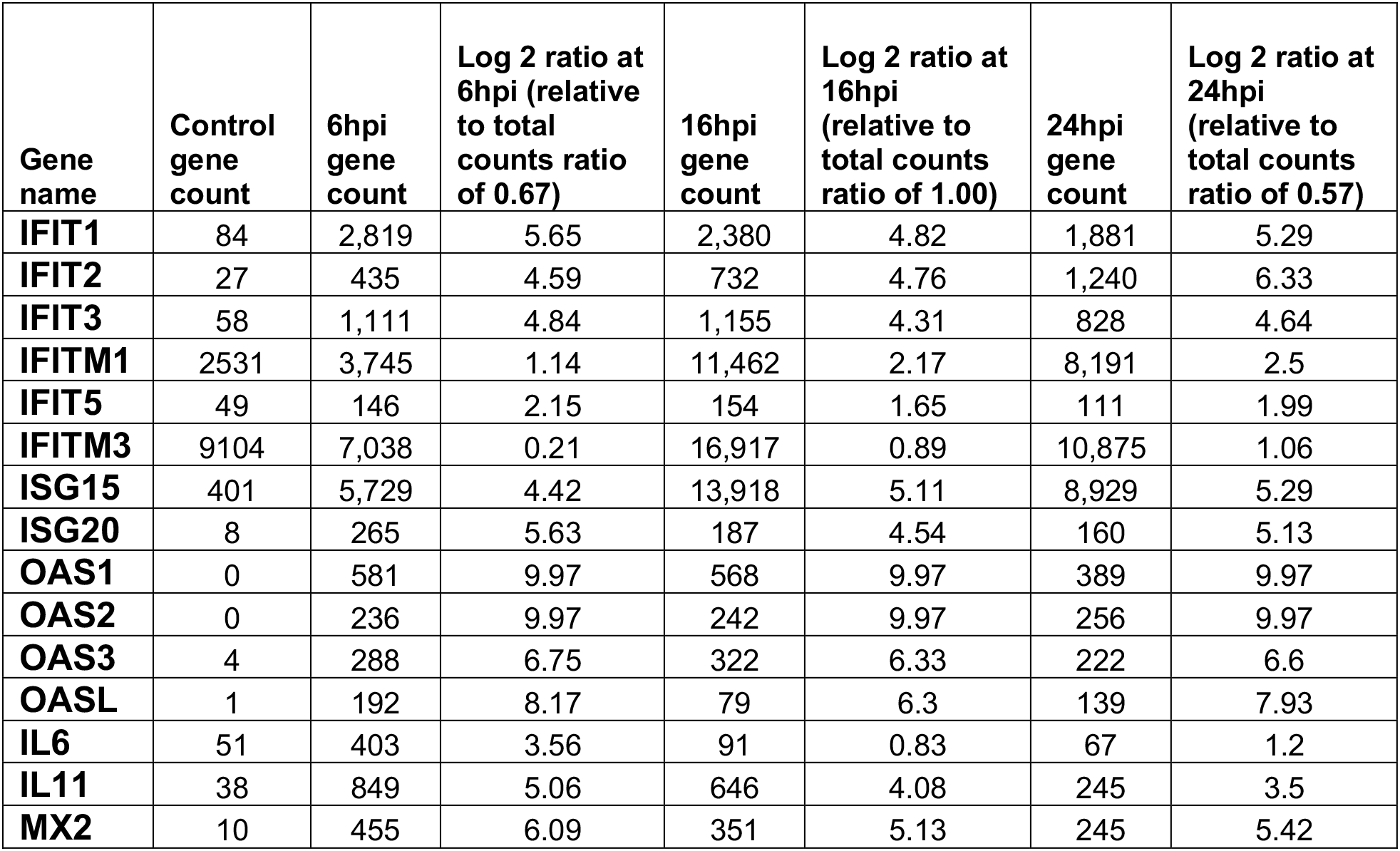
Selected human genes previously reported to show increased transcription post RSV infection. Table illustrating changes in numbers of detected transcripts at different time points post infection with RSV. The observed numbers of transcripts are corrected according to the total number of transcripts mapping to the human transcriptome at different time points and compared to the control sample. A log2 fold change is indicated - a ratio of 1, for example, indicates a doubling in the relative number of observed transcripts.

| Gene name | Control gene count | 6hpi gene count | Log 2 ratio at 6hpi (relative to total counts ratio of 0.67) | 16hpi gene count | Log 2 ratio at 16hpi (relative to total counts ratio of 1.00) | 24hpi gene count | Log 2 ratio at 24hpi (relative to total counts ratio of 0.57) |
| --- | --- | --- | --- | --- | --- | --- | --- |
| **IFIT1** | 84 | 2,819 | 5.65 | 2,380 | 4.82 | 1,881 | 5.29 |
| **IFIT2** | 27 | 435 | 4.59 | 732 | 4.76 | 1,240 | 6.33 |
| **IFIT3** | 58 | 1,111 | 4.84 | 1,155 | 4.31 | 828 | 4.64 |
| **IFITM1** | 2531 | 3,745 | 1.14 | 11,462 | 2.17 | 8,191 | 2.5 |
| **IFIT5** | 49 | 146 | 2.15 | 154 | 1.65 | 111 | 1.99 |
| **IFITM3** | 9104 | 7,038 | 0.21 | 16,917 | 0.89 | 10,875 | 1.06 |
| **ISG15** | 401 | 5,729 | 4.42 | 13,918 | 5.11 | 8,929 | 5.29 |
| **ISG20** | 8 | 265 | 5.63 | 187 | 4.54 | 160 | 5.13 |
| **OAS1** | 0 | 581 | 9.97 | 568 | 9.97 | 389 | 9.97 |
| **OAS2** | 0 | 236 | 9.97 | 242 | 9.97 | 256 | 9.97 |
| **OAS3** | 4 | 288 | 6.75 | 322 | 6.33 | 222 | 6.6 |
| **OASL** | 1 | 192 | 8.17 | 79 | 6.3 | 139 | 7.93 |
| **IL6** | 51 | 403 | 3.56 | 91 | 0.83 | 67 | 1.2 |
| **IL11** | 38 | 849 | 5.06 | 646 | 4.08 | 245 | 3.5 |
| **MX2** | 10 | 455 | 6.09 | 351 | 5.13 | 245 | 5.42 |

## Discussion

Our dRNAseq dataset provides a detailed analysis of RSV mRNA expression at three timepoints in MRC-5 cells, a genetically normal lung cell line. By using dRNAseq we can observe and quantitate viral mRNA without potentially confounding artefacts from, for example, reverse transcription or PCR amplification. In addition, because our analysis is confined only to transcripts for which we have quantitated the length of the polyA tail we can also be confident that we are observing and quantitating mRNA only, and not genomic/antigenome RNA or immature mRNA.

Our data shows a broad gradient of individual transcripts decreasing in abundance from the 3’ to the 5’ end of the genome. However, there is little detectable attenuation at some gene junctions, rather there are groups of genes with similar transcript levels, one for *NS1* the second comprising *NS2*, *N*, *P* and *M*, the third with *SH*, *G*, *F* and *M2* with *L* being the fourth. We noted a similar pattern at 6 hours post infection but with lower *N* transcript abundance than for P or M. In contrast to previous recent reports based on qRT-PCR (25) or our own prior Illumina RNAseq data (24) we do not see elevated levels of *G* protein transcript abundance. Instead, our dRNAseq data is more aligned with early RSV data based on slot blots and UV inactivation studies (16, 22). Whilst we do not see a strict correlation with those data, we do observe a general overall decline. We propose that discrepancies between our data and previously published Illumina or qRT-PCR based analyses likely result from reverse transcription and PCR based steps used in those techniques (but not in dRNAseq) introducing unintended bias in the quantification of transcripts. Indeed, a recent careful analysis of transcript abundance in related paramyxoviruses reported that for illumina based approaches at least, the techniques used to enrich for viral mRNA and sequence can affect abundance estimates with approaches that distinguish the strand sense of the Illumina based sequence data having a significant impact on transcript abundance estimation (42). This analysis highlighted the potential for genome and antigenome RNA to confound attempts to determine mRNA abundance since they too will be enriched by RNA complementarity in any polyA based enrichment strategy. Nonetheless it is notable that for several gene pairs the abundance does not decline at the gene junction, and in some cases, such as *N*-*P* and *G*-*F* there appears to be marginally more mRNA from the downstream gene. For the *P* and *N* transcripts this may be due to the *N* mRNA being longer (and therefore more prone to degradation) as dRNAseq is more likely to successfully read the full length of shorter mRNA molecules, and indeed the peaks for *N* and *P* sequence coverage are similar, indicating that this may well be the case. However, as *G* transcripts are shorter than *F* transcripts this argument does not explain the larger number of *F* transcripts. It is possible that *G* mRNA is less stable than *F* mRNA, accounting for the difference in these steady-state transcripts. This suggests that aspects of mRNA stability may also be a notable factor in determining the steady state levels of mature RSV mRNA species. It is also possible that there is some feature of G mRNA, such as secondary structure, that makes it more difficult to sequence so that it is unrepresented in the read depth analysis. There is also a formal possibility that on occasion the viral polymerase ignores a GS signal but is still able to progress to the subsequent GS and then re-initiate transcription. Evidence that this can occur has been found for a paramyxovirus, SV41, in which *M* transcripts are exclusively dicistronic *M-F* mRNAs, but monocistronic *F* transcripts are also produced, suggesting that some polymerase can scan from the *P* GE signal to the *F* GS signal, without transcribing *M(43)*.

Our data expands considerably on the production of polycistronic mRNA by RSV, showing for the first time the relative levels of different polycistronic messages produced by the viral polymerase. An early paper in which RSV transcripts were analysed by Northern blot analysis had revealed polycistronic transcripts containing two or three genes, but the relative levels of these transcripts were not determined (11). We find that in our data the *SH-G* intergenic appears to have the strongest GE signal and therefore the fewest readthrough events with the *NS1* GE signal being the weakest. These findings are supported by previous work (11, 24, 44). However, the significance of the strength of GE signals is unclear as previous work suggests that relatively high level of readthrough at the *NS1* and *NS2* GE signals does not play significant role in viral replication, either *in vitro* or *in vivo* (45).

In addition to the canonical transcripts corresponding to the viral genes, we also examined the levels of the M2-L transcript. The RSV M2-L gene junction is unusual in that the L GS signal lies upstream (rather than downstream) of the M2 GE signal. It was previously shown that the RSV polymerase can access the L GS signal by scanning backwards from the M2 GE signal following M2 mRNA release (18). However, the M2 GE signal has a similar termination efficiency as other RSV GE signals (46), and so if the polymerase simply responded to signals as it encounters them, it would be expected to release the L transcript at the M2 GE signal, primarily resulting in a 68 nt M2-L transcript (not including the poly A tail) and only producing full-length L mRNA as a readthrough transcript, similar to a polycistronic mRNA. However, the data presented here clearly show that L mRNA is the major transcript produced from the L GS signal. Even if only full-length transcripts were considered, the L mRNA was produced at a 5-6 fold excess over the M2-L transcript, and as noted in the results section, this is an under-representation of full-length L mRNA due to its large size. If we consider that the L mRNA 3’ end reads represent full-length L mRNAs then there was a 240-fold excess of L mRNA over M2-L transcripts at 24 hpi. An explanation for why the polymerase usually generates a full-length L mRNA versus the M2-L transcript comes from work with vesicular stomatitis virus, which showed that the polymerase can only recognize a GE signal positioned beyond a certain distance from the GS signal (47). This probably reflects the need for the polymerase to cap the mRNA before it can accurately recognize a GE signal (48). We also examined M2-L transcripts with a limited (<20 nt) poly A tail length, as in another study it was shown that VSV polymerase that does not cap the RNA can recognize a GE signal but fails to properly polyadenylate the mRNA (49). However, there was no evidence of such M2-L transcripts. Thus, we conclude that is probably insufficient distance before the M2 GE for it to be recognized by a polymerase that initiated at the L GS, allowing the polymerase to disregard this GE signal and synthesize full-length L mRNA, as suggested previously (47).

We observed in our datasets a number of unusual transcripts, some with minor deletions that could be a consequence of limitations with the nanopore sequencing technology. However, we also observed a substantial number of transcripts with more substantial deletions. In one example we detected the same 25 nt deletion in *NS2* transcripts in our datasets at both 16 and 24 hours post infection. In this example, given the number of distinct transcripts with the same deletion and the fact they were observed in two different samples suggests that these kinds of deletions are genuine and reasonably frequent. We cannot exclude the possibility that some rearrangements we detected could occur as an artefact of the sequencing technique, for example, at the DNA-RNA ligase step of sample preparation. However, previously we reported very rare splicing events in the adenovirus transcriptome and went on to show that even apparently rare transcripts (as few as 10 molecules in over 1.2 million reads) could be independently verified by directed RT-PCR (26) and our analysis of the adenovirus transcriptome suggests that there is a broad range of mRNA species made amongst a small number of highly dominant transcripts that correspond to the classical adenovirus transcriptome. This concept was subsequently independently verified for a different serotype of adenovirus, again using dRNA-seq (50). If the transcripts detected in the current study are genuine, it suggests the RSV polymerase has the potential to skip short distances when transcribing mRNA. Isolates of RSV have been found with small duplications in viral G gene, and defective interfering genomes are frequently generated, indicating that the viral polymerase can dissociate and reassociate with the template during genome replication (51–54). It is therefore not surprising to see a similar background of unusual mRNAs that do not match the classical transcripts in RSV infected cells.

We also noted the polyA tail length of the viral transcripts are similar to that reported for human transcript polyA tails (32) and in that respect unremarkable. To our knowledge this is the first time polyA tail length has been reported for RSV. Notably, these polyA tails are added by the viral polymerase but without any clear mechanism to explain how the length is determined or why they are broadly similar to host mRNA which is polyadenylated in the nucleus. In addition, between 16 and 24 hours there is a noticeable decline in polyA tail length across all the different mRNA classes. We have previously noted a decline in polyA tail length during adenovirus infections whose mRNA are made by host RNA pol II (26) and even earlier reports noted a similar effect during coronavirus replication (55). Potentially therefore a decline in polyA tail length may simply be a consequence of host cell function decline as infection progresses. Potentially of greater significance is the observation that transcripts with very short polyA tails are underrepresented for the *N*, *SH* and *G* genes (Figure 5 and Table 5). We retrospectively performed a similar analysis on our previously published work on SARS-CoV-2 transcripts (30) and did not observe any significant difference in the distribution of SARS-CoV-2 transcripts with very short polyA tails. That three RSV genes in two independent datasets (16 hpi and 24 hpi) show the same pattern of underrepresentation is notable and may reflect differences in rates of translation or mRNA stability both of which can be governed by polyA tail length (36). In principle the underrepresentation of *N*, *G* and *SH* may reflect higher rates of translation for those transcripts and experiments to examine this possibility are underway.

Our data also contains interesting information on gene expression of host cells. At this time there is no agreed consensus on the analysis of dRNAseq data which has a lower read depth than traditional Illumina based RNAseq methods. But our results do show that many genes which are known to increase on RSV infection do also show an increased expression in our time course (Table 6). Potentially further analysis of our data may provide additional insights into RSV modulation of host gene expression.

In conclusion, our dataset is the first dRNAseq transcriptomic analysis of RSV mRNAs produced over an infection time-course. This dataset provides fine detail of the transcriptomic repertoire of RSV in a cell culture system. Our findings align closely with early papers that attempted to determine the relative abundance of RSV transcripts and provides additional quantitative information that was lacking in the previous work. In addition, we believe that this analysis highlights the potential for inaccurate measurements when trying to estimate transcript abundance by indirect methods such as qRT-PCR or short-read based RNAseq techniques, and we think that this technical insight is broadly relevant to the study of viral gene transcription.

## Acknowledgements

We’d like to thank the BBSRC for funding (DAM and ID, grant BB/M02542X/1).

## Author Contributions

I.D-B., G.S., S.H., T.G., E.M. and J.L. carried out the laboratory work in this article. I.D-B, R.F., J. A.H. and D.A.M. conceived and designed the program of research, contributed intellectually to drafting the research text and critiqued the intellectual content. All authors contributed to the drafting of the manuscript. D.A.M. supervised the work, wrote the software and used it to analyse the data.

